# Social Isolation During Adolescence Differentially Affects Spatial Learning in Adult Male and Female Mice

**DOI:** 10.1101/2024.07.31.606027

**Authors:** Sadiyah Hanif, Mia Sclar, Jinah Lee, Caleb Nichols, Ekaterina Likhtik, Nesha S. Burghardt

## Abstract

Social isolation is a risk factor for cognitive impairment. Adolescents may be particularly vulnerable to these effects, because they are in a critical period of development marked by significant physical, hormonal, and social changes. However, it is unclear if the effects of social isolation on learning and memory are similar in both sexes or if they persist into adulthood after a period of recovery. We socially isolated male and female 129Sv/Ev mice throughout adolescence (post-natal days 29-56), provided a 2-week re-socialization recovery period, and then tested spatial learning and cognitive flexibility in the active place avoidance task. After behavioral testing, mice were injected with 5’-bromo-2’-deoxyuridine (BrdU) so that lasting effects of social isolation on cell proliferation in the dentate gyrus could be examined. We found that in males, isolation led to a modest impairment in the rate of initial spatial learning, whereas in females, initial learning was unaffected. However, when the location of the shock zone was switched during the conflict variant of the task, cognitive flexibility was impaired in females only. Similarly, social isolation reduced cell proliferation in the ventral dentate gyrus only in females. Together, these findings indicate that social isolation during adolescence differentially impairs spatial processing in males and females, with effects that persist into adulthood.

## Introduction

Humans are a social species whose cognitive function is negatively affected by social isolation. For example, perceived social isolation, or loneliness, has long-been known to be a risk factor for cognitive decline (Cacioppo and Hawkley 2009) and Alzheimer’s disease (Wilson et al. 2007). This was exemplified by the COVID-19 pandemic, during which social isolation resulting from the global lockdown significantly impaired cognitive performance in children and adults within the United States and at least 15 other countries (Ingram et al. 2021; Betthauser et al. 2023). Studies in students reveal that the pandemic not only had a detrimental effect on academic performance during the lockdown, but that students had not recovered a year after schools reopened (Breit et al. 2023; Di Pietro 2023; Fahle et al. 2023). It remains unclear how long-lasting these effects will be or the extent to which they are caused by the stress of social isolation alone.

Across species, adolescence is a critical period of development characterized by significant physical, hormonal, and social changes. It roughly corresponds to ages 10-24 in humans (Sawyer et al. 2018) and postnatal days (PND) 28-56 in rodents (Andersen 2003; McCormick and Mathews 2007). Given the increase in social behavior that occurs during this time (Thor and Holloway 1984; Blakemore and Mills 2014), adolescents may be particularly vulnerable to the detrimental effects of social isolation. In addition, sex hormones begin to regulate hypothalamic-pituitary-adrenal (HPA) function during adolescence (McCormick and Mathews 2007), potentially increasing the overall impact of stress during this period of development. Based on the dramatic differences between males and females in the hormonal changes that occur during puberty, it is possible that social isolation during adolescence affects each sex differently. The finding that there are sex differences in HPA axis activity in adulthood is in line with this possibility (Sencar-Cupovic and Milkovic 1976; Bangasser and Wiersielis 2018).

The hippocampus is a stress sensitive brain structure (McEwen et al. 2016) that continues to develop during adolescence (Spalding et al. 2013; Hueston et al. 2017) and is critical for learning and memory (Bird and Burgess 2008). Different types of social stressors have been shown to affect hippocampal-dependent learning. For example, social instability stress during adolescence impairs spatial memory in an object spatial location test (McCormick et al. 2010; McCormick et al. 2012) and maternal deprivation impairs spatial learning in the Morris water maze, but increases contextual fear conditioning (Oomen et al. 2010). Studies of social isolation stress during adolescence have primarily focused on spatial learning in the Morris water maze, but the results have led to contradictory findings. While some groups show that adolescent isolation impairs spatial learning and memory (Hellemans et al. 2004; Ibi et al. 2008), others show no effect (Schrijver et al. 2004; Voikar et al. 2005; Kulesskaya et al. 2011), or improvements in acquisition (Wongwitdecha and Marsden 1996).In addition, almost all social isolation studies were conducted in male rodents and sex differences have not been explored.

We tested the effects of socially isolating adolescent male and female 129Sv/Ev mice (PND 29-56) on active place avoidance, a spatial learning task that is sensitive to minor hippocampal dysfunction (Cimadevilla et al. 2001; Wesierska et al. 2005). During place avoidance, animals on a rotating arena learn to avoid a region of the room designated as a shock zone. In addition to evaluating spatial learning and memory, we used a conflict variant of the task to test cognitive flexibility, which we have previously shown to be dependent on dentate gyrus function (Kheirbek et al. 2013). We also examined effects of social isolation on cell proliferation in the dentate gyrus, as hippocampal neurogenesis undergoes dynamic changes during adolescence (Sousa et al. 1998; Spalding et al. 2013; Hueston et al. 2017; Boldrini et al. 2018) that could be altered by stress. Behavioral testing and cell quantification took place following a two-week recovery period so that the long-lasting effects of social isolation could be evaluated. We found sex differences in both the behavioral and cellular effects of social isolation, which included a modest impairment in spatial learning in males and impaired cognitive flexibility and reduced cell proliferation in the ventral dentate gyrus of females. This work highlights the importance of considering sex when investigating mechanisms mediating the long-term effects of social isolation and when developing interventions.

## Results

### Social isolation stress during adolescence does not have long-lasting effects on anxiety-like behavior

While the focus of our study was on spatial learning, social isolation has been widely reported to affect mood and anxiety in humans (Cacioppo et al. 2006; Leigh-Hunt et al. 2017) and animals (Fone and Porkess 2008; Lukkes et al. 2009). To investigate whether there are sex differences in the long-lasting effects of isolation on anxiety-like behavior, we used exploration of the novel rotating arena during pretraining as a modified open field test. A two-way ANOVA on total distance traveled revealed no sex x stress interaction (F_(1,29)_ = 0.17, p = 0.68) or main effects of sex (F_(1,29)_ = 7.29e-005, p = 0.99) or stress (F_(1,29)_ = 1.22, p = 0.28) (**Fig. 1A),** indicating that isolation stress did not affect exploration of the arena in either sex. Interestingly, a two-way ANOVA on the percent of total distance traveled within the center half of the arena revealed a significant effect of sex (F_(1,29)_ = 6.49, p = 0.02), with females walking in the center more than males (**Fig. 1B**). However, there was no sex x stress interaction (F_(1,29)_ = 1.72, p = 0.20) or main effect stress (F_(1,29)_ = 1.98, p = 0.17), indicating that adolescent stress did not affect anxiety-like behavior in adulthood in either sex when measured in this way.

**Figure 1.**
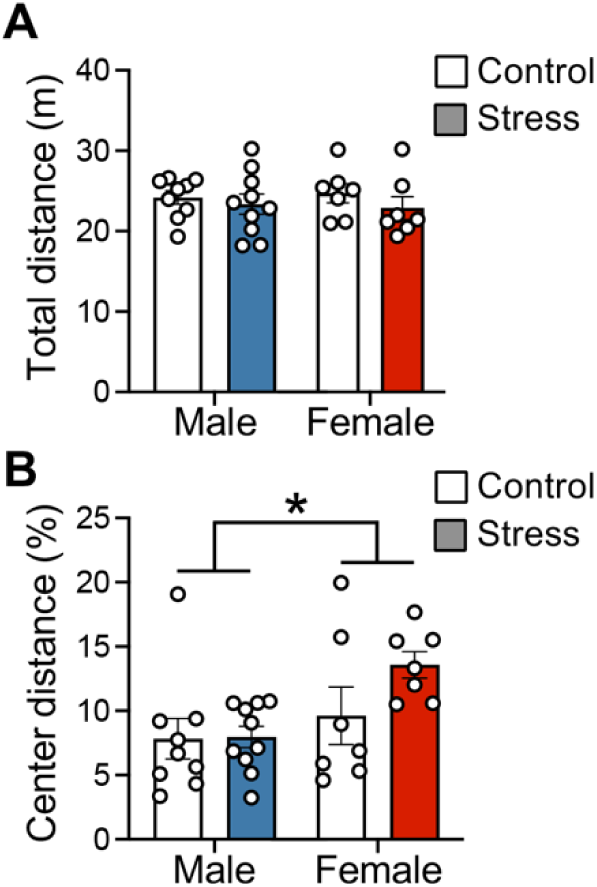
Social isolation during adolescence does not have long-term effects on exploration or anxiety-like behavior. **(A)** Total distance traveled on the rotating arena during pretraining. **(B)** Percent of total distance traveled in the center half of the rotating arena during pretraining. n = 9 control males; n = 10 stress males; n = 7 control females; n = 7 stress females. Data are represented as mean ± SEM. *p<0.05.

### Social isolation during adolescence leads to a modest impairment in the rate of spatial learning in males

We next investigated the long-term effects of social isolation on spatial learning in male mice using the active place avoidance paradigm (**Fig. 2A**). Before training began (pretraining), stressed males entered the inactive shock zone as often as non-stressed males (t_(17)_ = 0.23, p = 0.82) (**Fig. 2B, 2C**). Then during initial training, mice received a brief foot shock upon entering a stationary shock zone defined by cues within the room. Mice were trained to avoid this shock zone with three 10-min long trials a day for two consecutive days (Day 1: Trials 1-3; Day 2: Trials 4-6), during which the shock was always on and the shock zone was always in the same location. A two-way repeated measures ANOVA on number of entrances into the initial shock zone across all 6 trials revealed a main effect of trial (F_(3.87, 65.70)_ = 7.80, p<0.0001), indicating that performance improved with training. However, there was no trial x stress group interaction (F_(5,85)_ = 0.83, p = 0.53) (**Fig. 2B, 2C**). To evaluate whether stress affected retention of this spatial memory, we measured latency to first enter the shock zone at the beginning of the second day of training (trial 4), which was 24-hours after the last training trial. We found that stressed males entered the shock zone faster than controls (t_(17)_ = 2.09, p = 0.05) (**Fig. 2D**), indicating worse spatial recall. While this might reflect impaired memory consolidation during the 24-hour retention period, it could also be attributed to slower acquisition the previous day. To explore the latter possibility, we investigated learning during the first day of training (trials 1-3) more closely. We found that both groups learned to avoid the shock zone quickly, as demonstrated by an average of 2.56 (control group) and 3.0 (stressed group) entrances during the third 10-min training trial (trial 3). To better capture the rate at which learning occurred, we calculated the number of times mice entered the shock zone during the first half of each trial. While the repeated-measures ANOVA on entrances during the first 5 minutes of trials 1-3 revealed no significant stress x trial interaction (F_(2, 34)_ = 1.24, p = 0.30), planned comparisons of each trial with Bonferroni’s multiple comparisons test revealed that stressed animals entered the shock zone significantly more than controls on trial 2 (p = 0.006) (**Fig. 2E**). Similarly, a two-way repeated measures ANOVA on latency to first enter the shock zone on trials 2-3 revealed an effect of trial that approached significance (F_(1,17)_ = 3.91, p = 0.06), with Bonferroni’s multiple planned comparisons indicating that latency was lower in stressed males than controls on trial 2 (p = 0.05) but not trial 3 (p = 0.95) (**Fig, 2F**). Similarly, the time-in-location map illustrating where each group spent their time during each training trial shows subtle group differences during trials 1-2 (**Fig. 3A**). Collectively, these results indicate that by the end of the first trial, stressed males had not acquired spatial learning as well as controls, but were able to reach levels comparable to controls by the end of trial 3.

**Figure 2.**
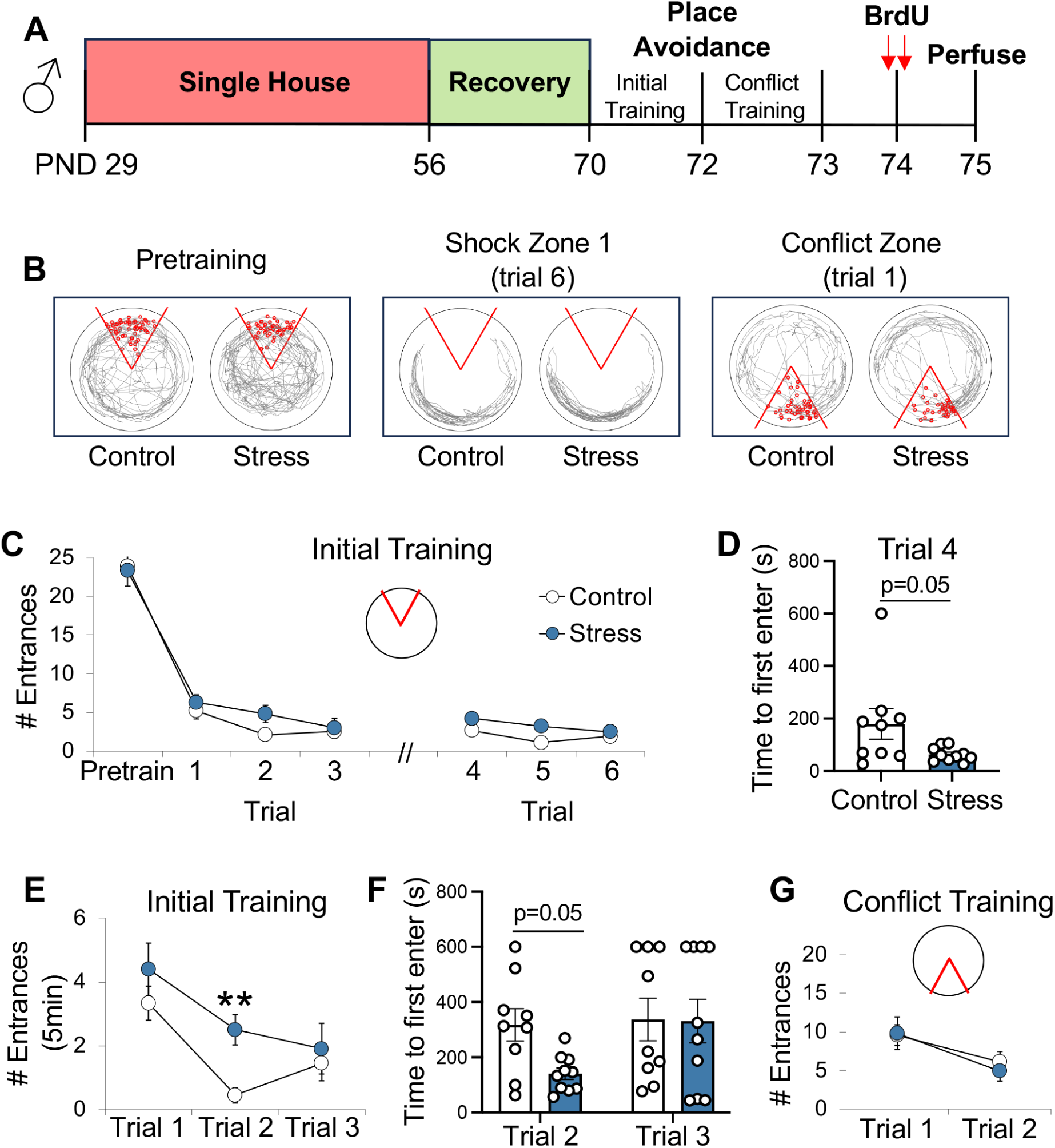
Social isolation during adolescence leads to a modest impairment in spatial learning in males. **(A)** Experimental timeline. **(B)** Tracked behavior (gray) of a representative control and stressed male mouse before the shock was turn on (pretraining), during the last trial of initial training (trial 6), and during the first trial of conflict training. With the exception of pretraining, red circles indicate where the animal was shocked during each trial. **(C)** Social isolation did not affect the total number of times male mice entered the shock zone during pretraining or initial training. **(D)** The latency to first enter the shock zone after 24-hours of retention was lower in stressed males than controls. **(E)** During initial training, stressed males entered the shock zone more than controls during the first 5-minutes of trial 2 and **(F)** entered the shock zone faster than controls on trial 2. **(G)** There were no group differences in the number of times males entered the new shock zone during conflict training. n = 9 control males; n = 10 stress males. Data are represented as mean ± SEM. **p<0.01 vs. trial 2 controls.

**Figure 3.**
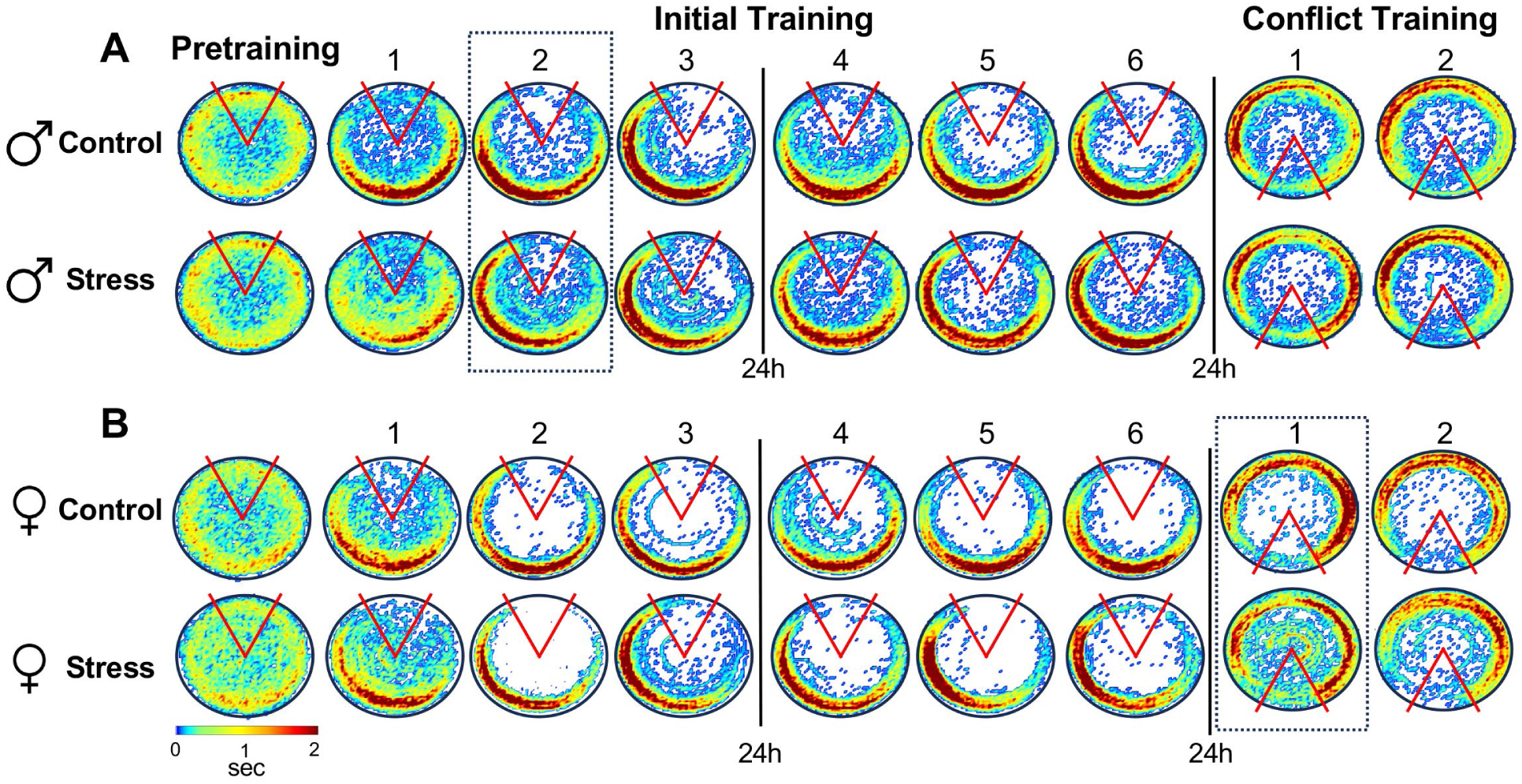
Social isolation affects patterns of avoidance behavior. Color-coded time-in-location maps for **(A)** males and **(B)** females of each group during pretraining, initial training (trials 1-6), and conflict training (trials 1-2). Colors represent group averages during each 10-minute trial, with blue indicating the lowest dwell time and red indicating the highest dwell time. The time represented by each color was the same for all maps (color bar, 0-2 sec). Dashed lines indicate trials during which stress increased number of entrances into the shock zone. n = 9 control males; n = 10 stress males; n = 7 control females; n = 7 stress females.

To evaluate cognitive flexibility, we used a conflict variant of the active place avoidance task, as previously described (Burghardt et al. 2012). This involved moving the shock zone 180° from its initial location, which challenged mice to flexibly adapt to a change in contingencies and suppress conditioned responses acquired during initial learning. A two-way repeated measures ANOVA on number of entrances into the new location of the shock zone across 2 conflict trials revealed a significant effect of trial (F_(1,17)_ = 21.31, p = 0002), but no trial x stress interaction (F_(1,17)_ = 0.65, p = 0.43) or main effect of stress (F_(1,17)_ = 0.06, p = 0.81) (**Fig. 2G**). Similarly, groups spent a similar amount of time within the new shock zone during the first conflict trial (t_(17)_ = 0.40, p = 0.69) (**Fig. 3A**). Together, these results indicate that social isolation during adolescence did not affect cognitive flexibility in adult males.

### Social isolation during adolescence impairs cognitive flexibility in females

To evaluate whether there are sex differences in the effects of social isolation on spatial learning, we tested female mice under the same conditions as males (**Fig. 4A**). During pretraining, we found that stressed females entered the inactive shock zone as often as female controls (t_(12)_ = 0.70, p = 0.50) (**Fig. 4B, 4C, 3B**). When the shock was turned on, a repeated measures ANOVA on number of entrances into the initial shock zone during all 6 trials revealed a significant effect of trial (F_(2.875, 34.49)_ = 4.44, p = 0.01), but no trial x stress interaction (F_(5, 60)_ = 0.55, p = 0.74) or main effect of stress (F_(1,12)_ = 0.33, p = 0.58) (**Fig 4C**). Both groups also exhibited a similar latency to first enter the shock zone on trial 4 (t_(12)_ = 1.02, p = 0.33) (**Fig. 4D**), indicating that retention of this spatial memory was not affected by isolation stress during adolescence. Furthermore, the repeated-measures ANOVA on number of entrances during the first 5 minutes of trials 1-3 revealed no significant stress x trial interaction (F_(2, 24)_ = 0.29, p = 0.75) and planned comparisons of each trial confirmed that there were no group differences on any trial (p>0.99). Despite being as successful as controls at avoiding the initial shock zone, the time-in-location maps showed that the stressed females were spending more time to the left of the shock zone than controls, indicating they learned to avoid the shock zone in a different way (**Fig. 3B, 4B**). Indeed, once the task was acquired (trials 2-6), stressed females spent significantly more time in this counterclockwise (CCW) position than controls (t_(12)_ = 2.22, p = 0.047) (**Fig. 4E**). This strategy was risky, because mice were closer to the shock zone on an arena that was rotating clockwise. If mice were inactive, they risked being rotated directly into that shock zone.

**Figure 4.**
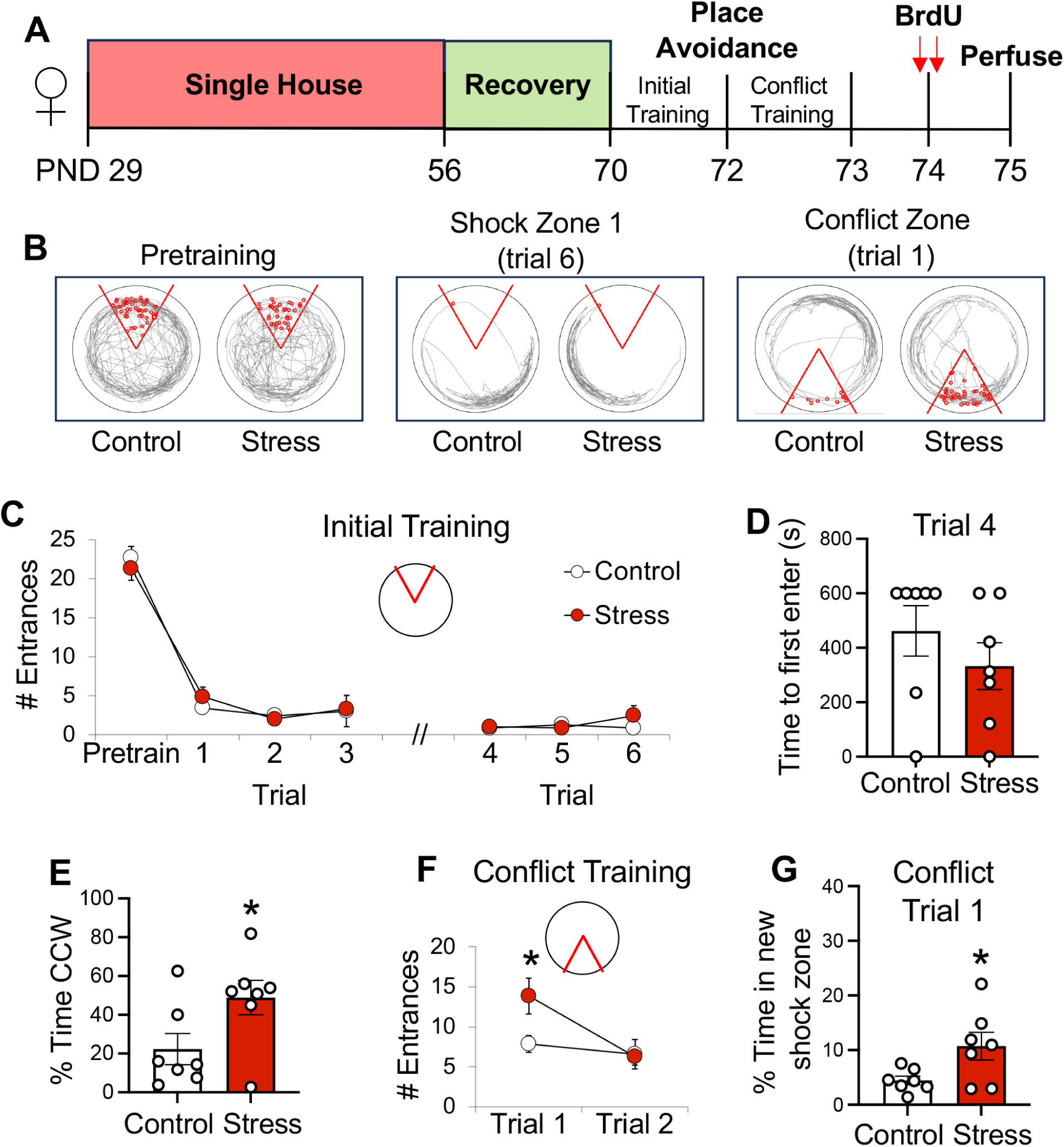
Social isolation during adolescence impairs cognitive flexibility in females. **(A)** Experimental timeline. **(B)** Tracked behavior (gray) of a representative control and stressed female mouse before the shock was turned on (pretraining), during the last trial of initial training (trial 6), and during the first trial of conflict training. With the exception of pretraining, red circles indicate where the animal was shocked during each trial. **(C)** Social isolation did not affect the number of times female mice entered the shock zone during initial training. **(D)** There were no group differences in latency to first enter the shock zone after 24-hours of retention. **(E)** During initial training, stressed females spent more time to left of the shock zone (i.e. counterclockwise, CCW) than controls. **(F-G)** During the first trial of conflict training, stressed females entered the new shock zone more and spent more time within that shock zone than controls. n = 7 per group. Data are represented as mean ± SEM. *p<0.05.

When the position of the shock zone was moved to the opposite side of the room, stressed females entered the new shock zone more than controls. A two-way repeated measures ANOVA on number of entrances during conflict training revealed a significant stress x trial interaction (F_(1,12)_ = 4.78, p = 0.049) and Bonferroni corrected post-hoc test confirmed that stressed females entered the new shock zone more than controls during the first conflict trial (p = 0.03) (**Fig. 4F**). Stressed females also spent more time within the new shock zone than controls during conflict trial 1 (t_(12)_ = 2.36, p = 0.036) (**Fig. 3B, 4G**), which is when flexibility was challenged the most. Together, these results demonstrate that unlike males, social isolation during adolescence in females impaired cognitive flexibility.

### Social isolation during adolescence reduces cell proliferation in females

We next evaluated the long-lasting effects of social isolation on cell proliferation in the hippocampus. Upon completion of behavioral testing, mice of both sexes were injected twice with 5’-bromo-2’-deoxyuridine (BrdU) and perfused the next day (**Fig. 2A, 4A**). Based on known functional differences between dorsal and ventral hippocampus (Fanselow and Dong 2010), we analyzed BrdU-positive (+) cells (**Fig. 5A**) in the dentate gyrus of each pole separately. In males, there was no effect of stress group (F_(1,11)_ = 0.17, p = 0.69), pole (F_(1,10)_ = 0.24, p=0.64) or stress x pole interaction (F_(1,10)_ = 1.67, p = 0.22), indicating that cell proliferation was not affected in this sex (**Fig. 5B**). In contrast, analysis of cell counts in females revealed a stress x pole interaction (F_(1,12)_ = 5.85, p = 0.03), with Bonferroni’s multiple comparisons test revealing a significant group difference in the ventral pole only (p = 0.036) (**Fig. 5A, 5C**). These results demonstrate a sex difference in the long-term effects of social isolation on cell proliferation and suggest that adult hippocampal neurogenesis may be reduced in the ventral hippocampus of females.

**Figure 5.**
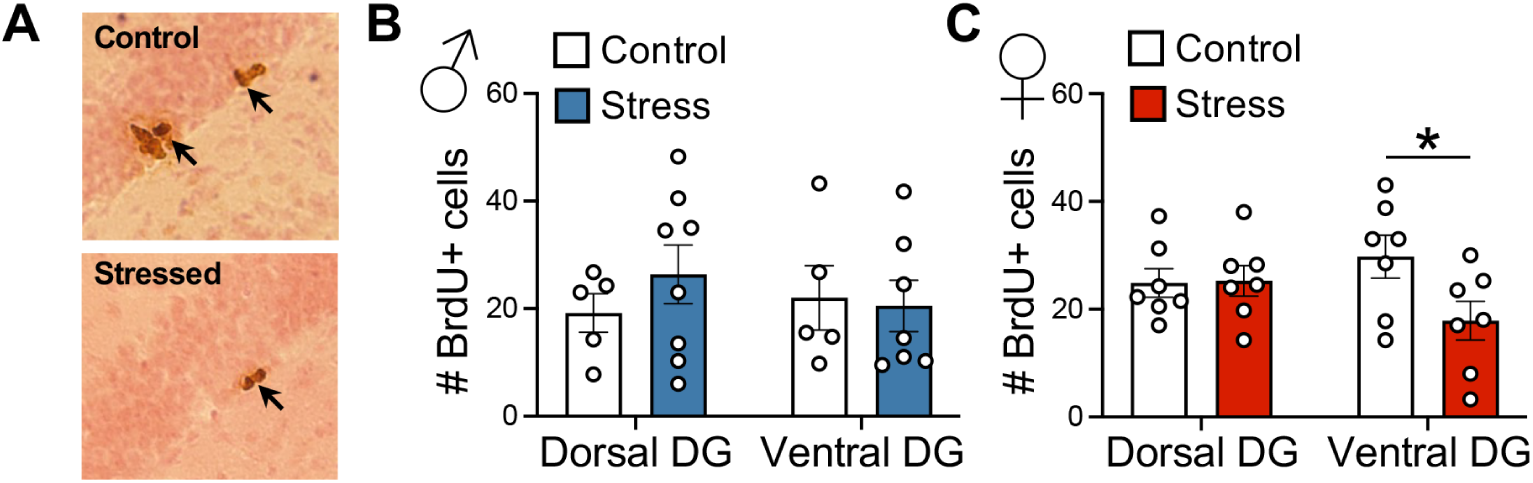
Social isolation decreases cell proliferation in females. **(A)** Representative images of BrdU immunostaining in the ventral DG of females in each group. Arrows indicate BrdU+ cells in the subgranular layer. **(B)** Social isolation did not affect the number of BrdU+ cells in the dorsal or ventral poles of males, but **(C)** decreased BrdU+ cells in the ventral pole of females. n = 7 control females, n = 7 stress females, n = 5 control males, n = 8 stress males. Data are represented as mean ± SEM. *p<0.05.

## Discussion

Using the active place avoidance paradigm, we tested the effects of social isolation throughout adolescence on hippocampal-dependent spatial learning in adulthood. We conducted our experiments in male and female 129Sv/Ev mice, a strain that exhibits high levels of anxiety-like behavior (Rodgers et al. 2002), is particularly vulnerable to social stress (Aubry et al. 2019), and is less aggressive than other commonly used strains (Abramov et al. 2008). This allowed us to re-house males during a recovery period with minimal concerns about fighting, so that both sexes could be tested under identical conditions. We found that social isolation led to a modest impairment in spatial learning in males, but impaired cognitive flexibility and reduced cell proliferation in the ventral dentate gyrus in females. Together, these results demonstrate that there are sex differences in the log-term effects of social isolation on learning.

Some previous studies have identified detrimental effects of social isolation during adolescence on performance in the Morris water maze (Hellemans et al. 2004; Ibi et al. 2008). Interestingly, the males tested in those studies were not given a recovery period, indicating that the modest deficit that we detect may represent a stress effect that is long-lasting rather than a delayed response to stress. However, numerous other studies in males of different species (rat or mice) report no effect of isolation on spatial learning in the Y-maze (Voikar et al. 2005) or water maze (Voikar et al. 2005; Han et al. 2011; Arakawa 2018). The discrepancy across findings could be attributed to a range of methodological differences, including variation in duration of isolation, mouse strain, or recovery period. Importantly, our study is the only one to asses spatial memory using the active place avoidance paradigm, which involves place responses that are less precise than those required by the water maze, but are more sensitive to learning impairments (Fenton and Bures 1993; Cimadevilla et al. 2001; Kubik and Fenton 2005; Serrano et al. 2008). Using active place avoidance may have therefore facilitated detection of the subtle impairment we found males.

While fewer studies have tested the effects of adolescent isolation on spatial learning in females, previous work indicates that it does not affect performance in the Y-maze or water maze (Einon and Morgan 1978; Kulesskaya et al. 2011). Similarly, we found that isolation did not affect the ability of females to avoid the initial shock zone location in place avoidance, but it did affect the strategy that was used. The pattern of avoidance behavior shown in the time-in-location map demonstrates that isolated females were less likely than controls to walk against the direction of rotation (clockwise) to identify the right edge of the shock zone, resulting in limited movement that was concentrated within the left-side of the room. As a result, stressed females were more at risk of being rotated into the shock zone than controls, who were directly opposite the shock zone. These findings may reflect an effect of social isolation on response to uncertainty and willingness to explore, such that isolated females were more reluctant to explore the right-side of the room and risk being shocked, than controls. Instead, stressed females remained in a part of the room that they already learned was safe, despite being close to the shock zone.

Cognitive flexibility requires adapting to a change in contingencies by modifying a learned response. Several studies have evaluated cognitive flexibility in socially isolated animals using tasks that are dependent on the prefrontal cortex (Makinodan et al. 2012; Baarendse et al. 2013; Hinton et al. 2019). Other studies have tested cognitive flexibility by using reversal learning in the water maze, with results indicating no effect (Voikar et al. 2005) or a partial impairment in isolated males (Han et al. 2011), and an impairment in isolated females in the absence of recovery (Kulesskaya et al. 2011). Here, we tested cognitive flexibility using a variant of the active place avoidance task, which we have previously shown to be sensitive to dentate gyrus function. In that work, optogenetically inhibiting the dentate gyrus or ablating adult neurogenesis from the dentate gyrus increased the number of times mice entered the new shock zone during conflict trials (Burghardt et al. 2012; Kheirbek et al. 2013). Interestingly, neither manipulation impaired the ability to learn the initial location of the shock zone, a form of learning that is more dependent on CA1 plasticity (Pavlowsky et al. 2017). The sex differences in behavior that we report here may therefore reflect sex-specific effects of social isolation on different hippocampal subregions. In males, the modest impairment in initial learning may be attributed to long-lasting effects of social isolation on CA1 functioning, while the deficit in cognitive flexibility in females may reflect stress-mediated changes in dentate gyrus function. Our finding that cell proliferation remained reduced in the ventral dentate gyrus of females is in line with this suggestion. There may also be additional stress-induced changes within mature granule cells of the dentate gyrus that contribute to the impairment in flexibility that we report.

Interestingly, the reduction in cell proliferation that we find in isolated females is specific to the ventral dentate gyrus. Similarly, social defeat, social submissiveness, maternal deprivation, and chronic unpredictable stress have all been reported to decrease adult neurogenesis in the ventral hippocampus only (Oomen et al. 2010; Anacker et al. 2018). Ventral hippocampus modulates HPA-axis function via projections to the bed nucleus of the stria terminalis (Anacker et al. 2011; Anacker 2014; Anacker et al. 2016) and suppression of adult hippocampal neurogenesis increases response of the HPA-axis (Schloesser et al. 2009; Snyder et al. 2011). In addition, silencing adult-born neurons in ventral dentate promotes stress susceptibility (Anacker et al. 2018), which together suggests that the stress-induced changes in ventral hippocampus that we report in females could affect subsequent responses to stress. Our finding that cell proliferation was intact in males indicates that females may be more vulnerable to these long-term effects of social isolation, potentially making them more susceptible to future stressors than males.

In conclusion, we tested the long-term effects of social isolation during adolescence in a stress-susceptible mouse strain. To our knowledge, this is the first study to test effects of social isolation on spatial learning and memory in both males and females so that sex differences could be identified. Unlike previous studies, which were primarily conducted with the Morris water maze, we tested mice in the active place avoidance paradigm, which is particularly sensitive to minor hippocampal dysfunction. Overall, we find that isolation affects females more than males. Specifically, isolated males exhibit a subtle impairment in spatial learning, while isolated females exhibit impaired cognitive flexibility and reduced cell proliferation in the ventral dentate gyrus. We suggest that these findings may reflect sex-specific effects of stress on different hippocampal subregions, potentially making females more vulnerable to stress later in life than males. This work demonstrates the importance of considering sex when evaluating the consequences of prolonged periods of isolation during adolescence on cognitive function.

## Materials and Methods

### Animals

129Sv/Ev mice were bred in-house using male and female breeders purchased at 8-weeks of age from Taconic Bioscience (Germantown, NY). Pups were weaned at postnatal (PND) 21, housed with littermates of the same sex, and maintained on a 12-hour light-dark schedule (09:00-21:00) with free access to food and water. All experiments were conducted in accordance with NIH-guidelines and approved by the Institutional Animal Care and Use Committee of Hunter College.

### Social Isolation

Mice were socially isolated throughout adolescence (PND 29-56) (Li et al. 2021) followed by a 2-week recovery period (Figure 2A, 3A), as outlined here. On PND 29, mice were either singly housed (stress group) or remained group-housed with littermates (control group) in cages containing a nestlet. Re-socialization began on PND 56, during which singly housed mice were housed together. Group-housed mice were also re-housed, but they were placed with novel cage mates, none of which were experimental mice. To minimize male aggression, which has been reported to increase with larger group size (Van Loo et al. 2001), males were always housed in groups of 2 and females were always housed in groups of 3, unless they were individually housed during isolation. During the 2-week recovery period, all mice were placed a new room in the animal facility in cages containing a nestlet and red igloo (Bio-Serve, Flemington, NJ).

### Active Place Avoidance

Active place avoidance training began on PND 70 and was conducted as previously described [Burghardt et al., 2012]. During a pretraining trial, mice walked freely on a circular platform (40cm diameter) that rotated clockwise (1 rpm), where they were exposed to visual cues within the room in the absence of shock. Then initial training began, during which mice received a brief constant current foot shock (500ms, 60Hz, 0.2mA, 1.5s inter-shock interval) when they entered a 60° stationary shock zone that was defined by cues within the room. An overhead camera connected to a tracking program (Tracker, Bio-Signal Group, Brooklyn, NY) recorded the position of the mouse and delivered shocks accordingly. Mice were given 3 initial training trials (Day 1: trials 1-3) followed by 3 additional initial training trials the next day (Day 2: trials 4-6) with the shock zone in the same position on each trial. Twenty-four hours later, conflict training began, during which the location of the shock zone was moved 180° from its initial position (Day 3: trials 1-2). This required mice to flexibly adapt to a change in contingencies. All trials (pretraining, initial training, conflict training) lasted 10 minutes, with an inter-trial interval of 50 minutes. The following behavioral responses were computed by Track Analysis software (Bio-Signal Group, Brooklyn, NY): number of times mice entered the shock zone, percentage of time in each quadrant of the arena, total distance traveled, and latency to first enter the shock zone.

### Time-In-Location Maps

Heat maps were generated as previously described (Burghardt et al. 2012). For each trial, the arena was divided into 0.64 cm^2^ pixels and time spent within each pixel was averaged across all animals in the same group. The location with the lowest dwell time (0.1 sec) is represented in blue and the highest dwell time (2 sec) is represented in red.

### BrdU Labeling

5’-bromo-2’-deoxyuridine (BrdU) (Roche, Indianapolis, IN) was dissolved in 0.9% NaCl and injected intraperitoneally (i.p.) twice at a dose of 75mg/kg, with 3 hours between injections (150mg/kg total). Twenty-four hours later, mice were deeply anesthetized with a mixture of ketamine (100 mg/kg, i.p.) and xylazine (7 mg/kg, i.p.) and transcardially perfused with cold PBS followed by cold 4% paraformaldehyde in PBS. Brains were removed, postfixed in 4% paraformaldehyde overnight, and then cryoprotected in 30% sucrose for at least 5 days in 4°C. Coronal sections (35μm) were cut through the hippocampus on a cryostat and stored in 0.1% NaN_3_. The day before immunohistochemistry began, sections were rinsed in PBS, mounted on slides, and left to dry at room temperature. The next day, slides were immersed in 10 mM citrate buffer (pH 6.0) for 2 hours at 95°C, rinsed in PBS, and then incubated in primary antibody (mouse anti-BrdU, 1:100, BD Biosciences, product number 555627) in 0.1% Triton X-100 overnight at room temperature. Slides were then rinsed in PBS, incubated in biotinylated secondary antibody (goat anti-mouse; 1:100, Jackson ImmunoResearch) in 0.1% Triton X-100 for 1 hour, treated with avidin-biotin-peroxidase complex (ABC Elite Kit, Vector Laboratories), stained with 3,3’ diaminobenzidine, and counterstained with Nuclear Fast Red (Vector Laboratories). BrdU+ cells in the granule cell layer of the dentate gyrus were quantified manually on an Olympus DP73 microscope by an investigator blind to group. Cells were counted bilaterally from sections containing dorsal hippocampus (bregma −1.34 to −2.03mm) and ventral hippocampus (bregma −2.94 to −3.51mm) using both the 20x and 40x objectives.

### Statistical Analysis

Data were analyzed with Student’s *t* test for independent samples, mixed effects modelling, or an ANOVA with Bonferroni post-hoc correction. All analyses were conducted using GraphPad Prism software (GraphPad, San Diego, CA).

## Acknowledgments

We thank Dr. Patricia Glennon, Barbara Wolin, and Sonia Acevedo for expert animal care. This work was supported by National Institutes of Health (NIH) grants R21 MH114182 (N.S.B. & E.L.), R21 MH135430 (N.S.B & E.L.), R01 MH118441 (E.L.), and PSC-CUNY Awards (N.S.B).

